# The length distribution and multiple specificity of naturally presented HLA-I ligands

**DOI:** 10.1101/335661

**Authors:** David Gfeller, Philippe Guillaume, Justine Michaux, Hui-Song Pak, Roy T. Daniel, Julien Racle, George Coukos, Michal Bassani-Sternberg

**Affiliations:** Department of Oncology UNIL CHUV, Ludwig Institute for Cancer Research, University of Lausanne, Switzerland.; Swiss Institute of Bioinformatics (SIB), Lausanne, Switzerland.; Department of Oncology UNIL CHUV, Ludwig Institute for Cancer Research, University Hospital of Lausanne, Switzerland.; Service of Neurosurgery, University Hospital of Lausanne, Switzerland

**Keywords:** HLA-I ligand predictions, Antigen Presentation/Processing, HLA peptidomics, Computational Immunology

## Abstract

HLA-I molecules bind short peptides and present them for recognition by CD8+ T cells. The length of HLA-I ligands typically ranges from 8 to 12 amino acids, but variability is observed across different HLA-I alleles. Here we collected recent in-depth HLA peptidomics data, including 12 newly generated HLA peptidomes (31,896 unique peptides) from human meningioma samples, to analyze the peptide length distribution and multiple specificity across 84 different HLA-I alleles. We observed a clear clustering of HLA-I alleles with distinct peptide length distributions, which enabled us to study the structural basis of peptide length distributions and predict peptide length distributions from HLA-I sequences. We further identified multiple specificity in several HLA-I molecules and validated these observations with binding assays. Explicitly modeling peptide length distribution and multiple specificity improved predictions of naturally presented HLA-I ligands, as demonstrated in an independent benchmarking based on the new human meningioma samples.

## Introduction

The binding of short intracellular peptides to Human Leukocyte Antigen Class I (HLA-I) molecules is a key step for immune recognition and prediction of naturally presented HLA-I ligands is currently the most important and most selective feature used to predict T cell epitopes. Peptide HLA-I interactions have been extensively studied either with binding assays or by Mass Spectrometry (MS). In recent years, improvements in sensitivity of MS instruments has led to the identification of hundreds of thousands naturally presented peptides, which include cancer testis antigens and neo-antigens (1, 2). Computational tools developed by ourselves and others have shown that HLA-I restriction can be predicted in large HLA-I peptidomics datasets, even in the absence of *a priori* information about HLA-I binding specificity using motif deconvolution (3, 4) or peptide clustering approaches (1, 5, 6). The use of MS data combined with these novel algorithms has led to refinement of known HLA-I motifs, identification of novel motifs, characterization of N- or C-terminal extensions and improved HLA-I ligand predictions (3, 4, 7–15).

In addition, HLA peptidomics data provide information about the length distribution of naturally presented HLA-I ligands. Interestingly, in a recent study, clear discrepancies were observed between length distributions of eluted peptides and length distributions predicted based on binding affinity for five soluble HLA-I alleles transfected into HeLa cells (16). This work indicated that the peptide length distribution observed in naturally presented HLA-I ligands is the result of both distinct affinity values of HLA-I alleles for peptides of different lengths as well as length biases in the pool of peptides available for loading onto HLA-I molecules in the Endoplasmic Reticulum (ER). However, most studies investigating peptide length distributions have been restricted to a few alleles (2, 5, 7, 8, 16, 17). Moreover, most existing HLA-I predictors, with the exception of recent versions of NetMHCpan (17) and NetMHC (18), do not include length distribution corrections based on MS data.

Another important feature characterizing peptides interacting with HLA-I molecules is the potential influence of a given amino acid at one position on the amino acid preferences at other positions. This type of co-operative effects, which often reflect multiple specificity patterns (19), can be captured with neural networks, support vector machines or mixture models. Such models have been widely used to predict the binding of peptides to HLA-I molecules (18, 20) or to peptide recognition domains (19, 21, 22). However few examples of actual amino acid correlations and multiple specificity have been documented among HLA-I ligands (3, 23). It remains therefore unclear how prevalent these correlations are in HLA-I alleles.

In this work, we set out to analyze the peptide length distribution and multiple specificity of more than 80 HLA-I molecules using high quality HLA peptidomics data generated by ourselves and others. To this end, we expanded our recent motif deconvolution algorithm MixMHCp (3) to better handle peptides of different lengths. Clear clustering of HLA-I alleles based on their peptide length distribution similarity was observed, which enabled us to investigate the structural basis of peptide length distributions and predict peptide length distributions based on HLA-I sequences. We then used MixMHCp to investigate multiple specificity in HLA-I molecules and validated these observations with binding assays. Both peptide length distribution and multiple specificity were then incorporated into our HLA-I ligand predictor MixMHCpred (4), leading to improved prediction accuracy of naturally presented peptides in ten meningioma samples. Finally, this work expands significantly the allelic coverage of MixMHCpred (122 HLA-I alleles in total), making it suitable to run predictions of naturally presented HLA-I ligands for most patients of caucasian origin.

## Materials and Methods

### Expanding motif deconvolution to peptides of different lengths

Binding motif deconvolution in HLA-I peptidomes with the previous version of the mixture model (MixMHCp1.0) performs well in many samples for 9- and 10-mers (3, 4). However, it often fails for 8-mers and for longer (>10-mers) peptides, since much less data is available for them in MS samples. To overcome this limitation, we introduce here a new approach to better handle peptides of different lengths. Motif deconvolution is first carried out on 9-mers as previously done in MixMHCp1.0 (3, 4). For peptides of other lengths, the motifs used in the mixture model are fixed, based on the first three and last two positions of the 9-mer motifs, and only the weights of the different motifs are learned from the data. The underlying justification for this approach is that HLA-I motifs, and especially positions close to the anchor residues, differ only slightly between 9-mers and other peptide lengths. However, the proportion of peptides from each allele in HLA-I peptidomics samples is likely to differ substantially for different peptide lengths, because of the different length distributions of HLA-I alleles. In addition, we integrated a flat motif (fixed values at all positions) in the mixture model to which peptides that do not match any motif inferred from the data can be assigned. This is useful to remove potential contaminants in MS data (6) and the number of peptides assigned to the flat motif is listed for each sample analyzed in this work in Supplemental Table I (see also Fig. 1). In mathematical terms, the new log-likelihood function for the peptides of length equal to core length (hereafter core length equals to 9) is defined as:

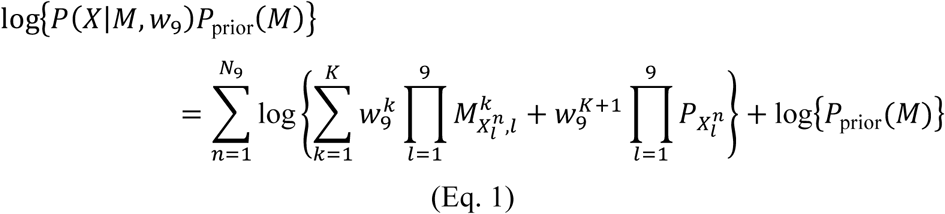

where *M*^*k*^ represents the *k*^th^ motif, 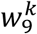 represents the weight of this motif in the 9-mer peptides, *N*_9_, the number of 9-mer ligands, 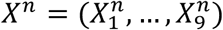 the n^th^ ligand of length 9 and *P*_prior_(*M*) the prior on the PWM entries as defined in (19, 22). The *P*_*i*_ *(i=1,…,20*) values are taken as the amino acid frequencies in the human proteome and represent a flat motif to which peptides that do not match any of the other motifs can be assigned, similar to the trash cluster in GibbsCluster (5, 24). This log-likelihood can be efficiently maximized used Expectation-Maximization (19, 22). For 9-mers, the only difference with the previous version of MixMHCp comes from the inclusion of the flat motif.

**FIGURE 1:**
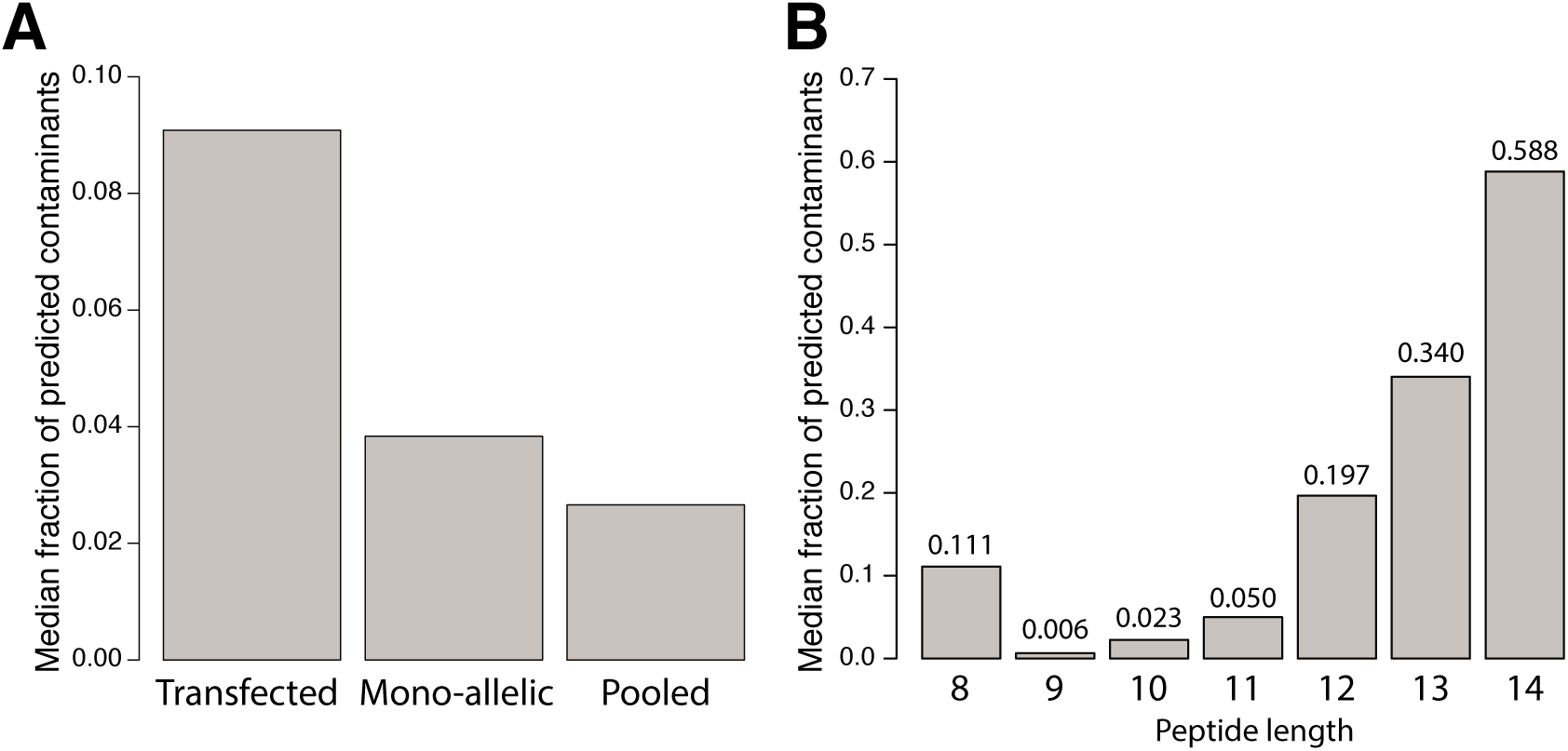
**A:** Median fraction of predicted contaminants in HLA peptidomics studies based on transfected soluble alleles, mono-allelic cell lines or pooled samples. **B:** Median fraction of contaminants among eluted peptides of different lengths.

For peptides of length L<9, the log-likelihood function reads:

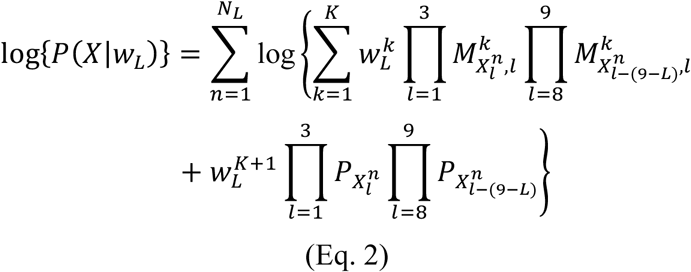

In this case the matrices *M*^*k*^ are fixed based on the motifs deconvoluted in the 9-mer peptides and only the weights of the different motifs are learned from the data, using again the Expectation-Maximization algorithm. For example, for 8-mers, it means that position 1-3 of the 9-mer motifs are used to score position 1-3 of the peptides, and position 8 and 9 of the 9-mer motifs are used to score position 7 and 8 of the peptides. For peptides of length L>9, the log-likelihood function is given by:

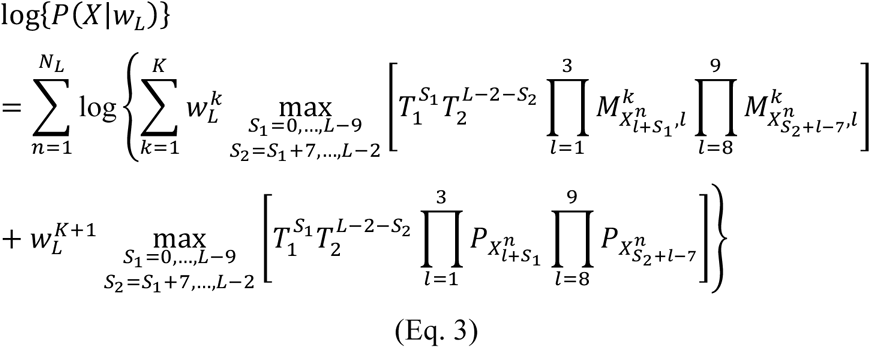

Here again the motifs themselves (*M*^*k*^*, k*=1-3, 8-9) are fixed based on those inferred from the 9-mer peptides and the weights 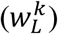 of the different motifs are learned separately for each length. The use of the ‘max’ function is meant to consider all possible cases of N- and C-terminal extensions (9). Information about which starting position of the N-terminal motif (i.e., S_1_+1) and end position of the C-terminal motif (i.e., S_2_+2) gave rise to the highest value for each cluster is given in output of MixMHCp2.0 (responsibility files), thereby providing predictions about potential N-and C-terminal extensions. *T*_*1*_=0.05 and *T*_*2*_=0.2 correspond to penalties assigned to extensions and represent the fact that, on average, cases where a motif can be found both at the terminus and within the peptide sequence correspond to bulges (9). The optimal number of motifs and the assignment of the different motifs to different alleles in each sample was determined as described in our previous work based on 9-mers (3, 4) and all motif annotations were manually reviewed.

The new version of the motif deconvolution (MixMHCp2.0) is available at https://github.com/GfellerLab/MixMHCp. All logos were generated with the LOLA software (http://baderlab.org/Software/LOLA).

### Collection of publicly available HLA peptidomics data

Publicly available HLA peptidomics datasets from mono-allelic studies (7, 8, 25, 26) and pooled studies (1, 2, 4, 14, 27–30) were considered in this work (Supplemental Table II). Almost all these samples were measured with 1% FDR and none of them was filtered by existing predictors. For the samples from (29), raw MS data were reprocessed and the list of peptides is available in (4). For the samples from (8), the list of peptides before filtering was kindly provided to us by the authors of this study. Finally IEDB (31) data (as of March 2018) were also collected, treating separately those annotated as “ligand presentation” (MS data) and those not (non-MS data).

### Motif deconvolution in HLA peptidomes

All HLA peptidomics datasets were analyzed with the new version of MixMHCp (see above) and all motifs were manually verified. These include mono-allelic samples, where the flat cluster was used to remove potential contaminants and motifs from endogenous alleles in samples transfected with soluble alleles (e.g., HLA-B35:03 and HLA-C04:01 in the data from (8)). Peptides were assigned to specific alleles based on the highest responsibility value observed for the corresponding motif (3, 4). For each allele, a unique list of peptides was derived by pooling together all peptides corresponding to motifs annotated to this allele across all samples, as previously described (3, 4). d. Peptides assigned to the flat motif in any sample were also excluded from downstream analyses.

### Length distribution in HLA-I ligands

The number of ligands *N*^*h*^(*L*) of each length *L*=*8,…,14* for a given allele *h* was computed based on available MS data. When MS data from our curated set of HLA peptidomics studies were not available for a given allele, we used MS data from IEDB. Although allele restriction was often predicted with former version of HLA-I ligand predictors in IEDB MS data, the utility of such data to estimate peptide length distributions and improve predictors has been recently demonstrated (17). Only HLA-I alleles with at least 200 peptides were considered in this analysis (84 alleles in total). Low-dimensionality representation of the alleles based on their peptide length distribution similarity was carried out with standard t-SNE, as implemented in R (initial_dims=5 and perplexity=28) (32). Of note the exact positioning of the points in the t-SNE plot was not used in downstream analyses, and is only meant to help visualization. Clustering of HLA-I alleles was performed in R (hclust package in R) based on the Euclidean distance between length distributions for all pairs of alleles (k=3, method=“ward.D2”). The average length distribution of each cluster was then computed (Fig. 2B).

**FIGURE 2:**
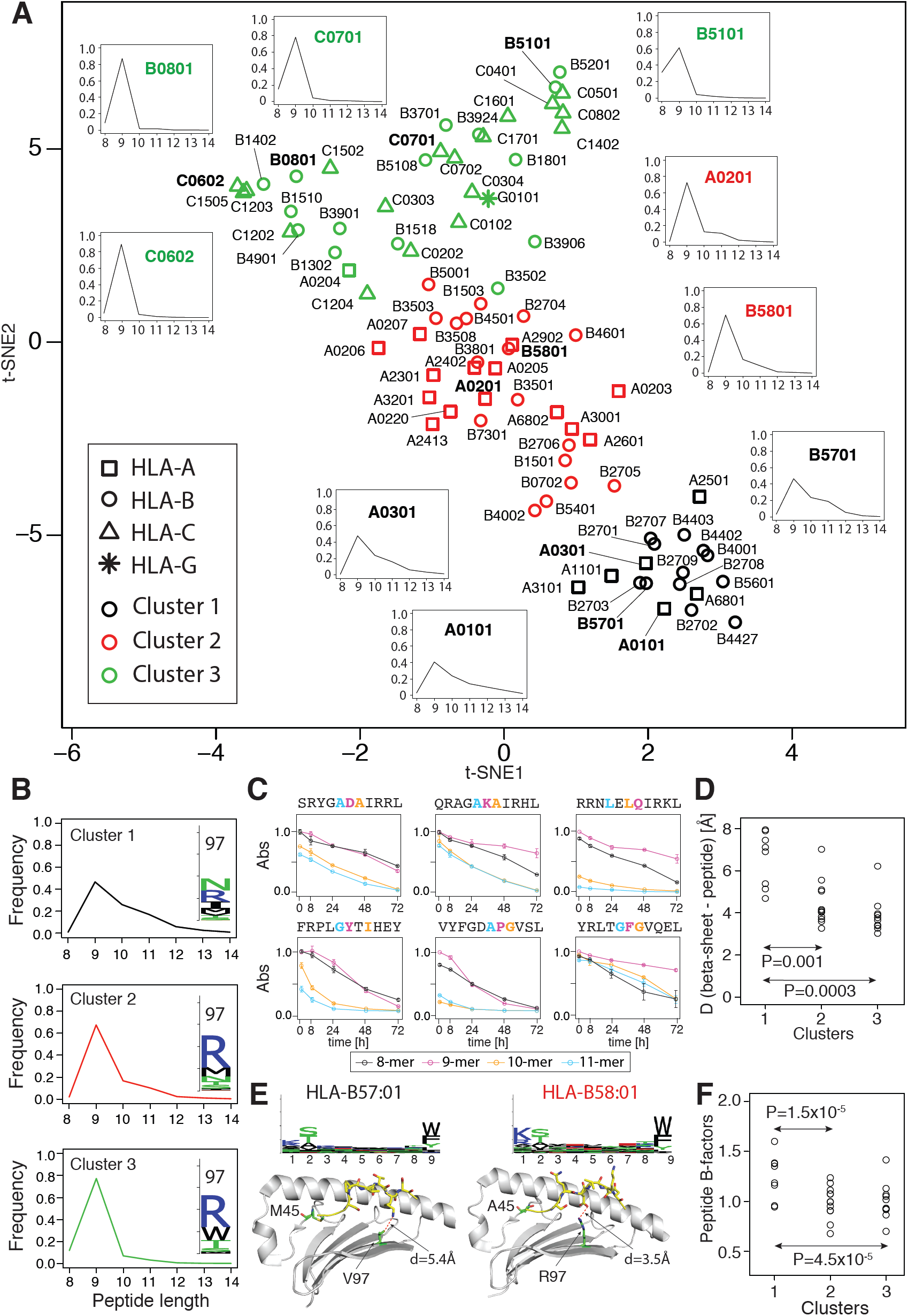
**A:** Low dimensionality representation (t-SNE) of HLA-I alleles with available MS data based on their peptide length distribution similarity. Different colors stand for different clusters and different shapes indicate HLA-A/-B/-C or -G alleles. **B:** Average peptide length distributions in each cluster. The logos shown in inset display the amino acid distribution at position 97 in the HLA-I alleles of each cluster. **C:** Binding stability of 8-, 9-, 10- and 11-mer peptides in complex with HLA-C06:02 obtained by ELISA (all values are renormalized by the 9-mer at t=0h). Residues shown in colors indicate those added in 9-mers (purple), 10-mers (orange) and 11-mers (cyan). Y-axis values correspond to ELISA absorbance signal normalized the signal for 9-mers at time t=0h. **D:** Distribution of the smallest distances between residues in the beta sheet and residues at P3-P8 in the peptide for alleles with available X-ray structures in each cluster (distances are measured between heavy atoms). **E:** Comparison of HLA-B57:01 and HLA B58:01 binding motifs and X-ray structures (PDB: 2RFX for HLA-B57:01 (35) and PDB: 5IND for HLA-B58:01 (36)). The two non-conserved residues in the HLA-I binding site are highlighted in green (M45 and V97 for HLA-B57:01 and A45 and R97 for HLA-B58:01). Peptide residues (P3 to P8) are displayed with yellow sticks. The smallest distance between residue 97 and the peptide (P3 to P8) is displayed in red. P-values correspond to Mann-Whitney *U* test. **F:** Normalized average B-factors of C*α* atoms at P3-P8 in the peptide for alleles with available X-ray structures in each cluster.

When incorporating peptide length distributions into our predictor (i.e., to compute the shifts, see below), a smoothing procedure was applied based on the average distribution 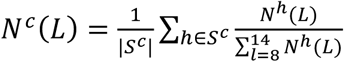 for each cluster *c* (*c=1,2,3*):

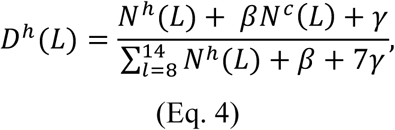

with *β*=100, *γ*=10 and *c=1, 2* or *3* depending on the cluster of allele *h* and *S*^*c*^ the set of alleles in cluster *c*. This will be especially useful for alleles without MS data (*N*^*h*^(*L*) = 0), see below.

### Prediction of peptide length distributions

To predict peptide length distribution directly from HLA-I sequences, we aligned the amino acid sequences of all alleles with at least 200 ligands found by MS. We then trained a multinomial logistic regression to predict the clusters observed in Fig. 2A (*cv.glmnet*, as implemented in R), using as input the amino acids at each position in the HLA-I binding site (i.e., positions 5, 7, 9, 33, 45, 59, 63, 66, 67, 69, 70, 73, 76, 77, 80, 81, 84, 97, 99, 114, 116, 123, 142, 143, 146, 147, 152, 156, 159, 163, 167 and 171, following residue numbering in X-ray structures), based on standard 20-dimensional encoding of amino acids. Cross-validation was carried out by randomly splitting into five groups the alleles shown in Fig. 2A and iteratively training the model (including both the clustering and the logistic regression) on four groups and testing the predictions on the 5^th^ group. In the testing set, the predicted length distribution of a given allele was taken as the average length distribution over the alleles of the cluster to which this allele was assigned and the correlation between the predicted length distribution and the actual one was computed. Average correlation reported in the manuscript corresponds to the average over all alleles and over ten different random choices of the groups used in the cross-validation (distinct random seeds). For comparison, correlations obtained by random cluster assignment of alleles in the testing sets were computed. This is important since peptide length distributions of all HLA-I molecules share common features, so we always expect some level of correlation between predicted and actual peptide length distributions, even with a random predictor.

Finally, for HLA-I alleles without MS data, the clusters were predicted with the logistic regression trained on the full dataset of alleles with MS data (84 alleles in total) and the predicted peptide length distributions for these alleles correspond to the average length distribution of the alleles belonging to the cluster where it was assigned, plus a flat pseudo-count given by *γ* (Eq. 4).

### Amino acid frequencies at non-anchor positions

Alleles with at least 20 ligands for all lengths *L=8,…14* were selected (i.e., A01:01, A02:01, A03:01, A11:01, A24:02, A29:02, A31:01, A68:01, B07:02, B15:01, B27:05, B44:02, B57:01) and the frequency of amino acids (*fr(x), x=1,…20*) at positions P5 to P Ω −2 was computed for each peptide length and each allele. For each amino acid, the average frequency (<*fr(x)*>) divided by the amino acid frequency in the human proteome (*P*_*x*_) and standard deviation across the 20 alleles are shown in Fig. 3. Amino acids have been ranked based on the slope of the linear regression (red line in Fig. 3). P-values in Fig. 3 correspond to the P-value of the Pearson correlation.

**FIGURE 3:**
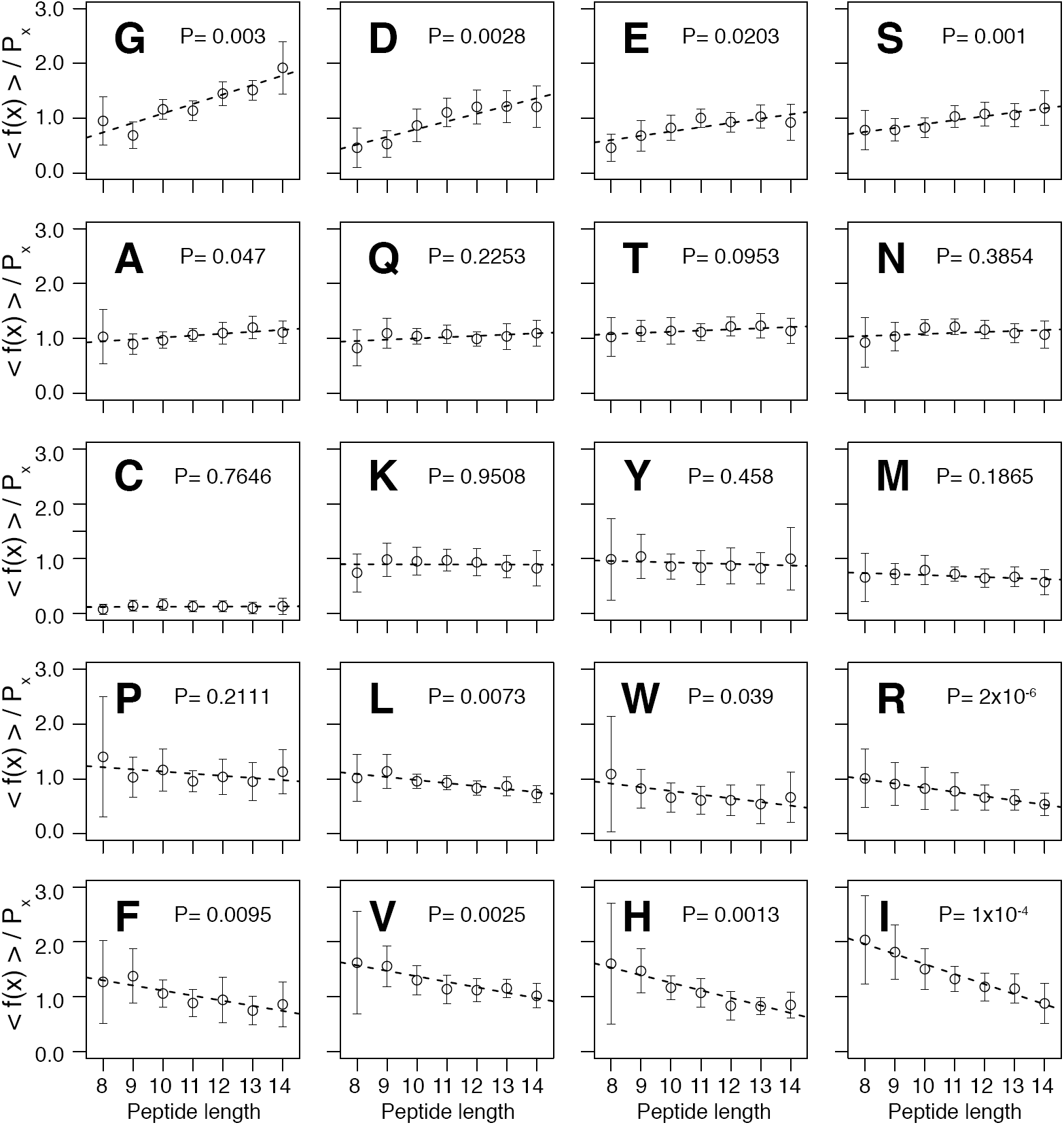
Average amino acid frequencies <*f(x)>* at non-anchor positions (P5 – P Ω-2) divided by the frequency in the human proteome (*P*_*x*_) as a function of peptide length. Each point represents the average over all alleles with at least 20 peptides for each length. Error bars represent standard deviations across these alleles. The black line corresponds to the linear fit. P-values correspond to the p-value of the correlation coefficient. Graphs have been ordered based on the slope of the black line.

### Building Position Weight Matrices

The training set of our HLA-I ligand predictor MixMHCpred was built as previously described (4), adding two new meningioma samples measured in this work (3911-ME_I and 4021_I, representing 4,719 unique peptides of length L=8,…,14) and several recent HLA peptidomics studies (8, 25, 26, 30). For alleles with less than 200 peptides, MS data from IEDB (31) were further included. For alleles that still had less than 20 peptides, *in vitro* binding data from IEDB were used (treating as positives all entries not annotated as “Negatives”). This led to a total of 122 alleles for which we could obtain ligands and build PWMs for length *L*=8 to *L*=14. To this end, all peptides assigned to the same allele across all studies were pooled together and each peptide was considered only once. Single Position Weight Matrices for each allele *h* and each peptide length *L* (*M*^(*h,L*)^) were built, including pseudo-counts based on BLOSUM correction and renormalization by the expected background amino acid frequencies (4, 33). In this framework, the score of a *L*-mer peptide *X* = (*X*_1_, …, *X*_L_) with allele *h* was computed as: 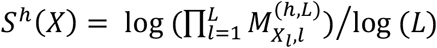. In a few cases, no ligand was available for L=8, and we used the 9-mer PWMs, removing the middle position with the lowest binding specificity. In a few other cases, ligands were not available for longer peptides (*L*=10,…, 14). In these cases, a non-specific position was added to *M*^(*h,L–1*)^ right after position *L*-4. Of note, these cases contribute only marginally to the predictions when incorporating peptide length distribution in the predictor, since by construction the corresponding lengths are given a very low weight (see below).

### Explicitly modeling length distribution in HLA-I ligand predictors

The scores given by standard PWMs result in peptide length distributions that do not follow those observed in MS data. To explicitly model the peptide length distributions of naturally presented ligands, we added a correction factor to the scoring of peptides of different lengths:

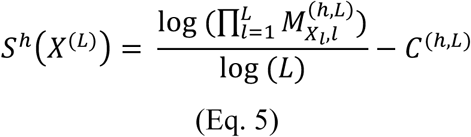

The correction factors *C*^(*h,L*)^ were computed as the score corresponding to 100 ×*D*^*h*^(*L*) percentile rank for 100’000 *L*-mers (randomly selected from the human proteome) without the correction factor. In this way, the top 1% peptides predicted from a set of 700’000 8- to 14-mers (100’000 of each length) follows by construction the distribution *D*^*h*^(*L*) derived from MS data and positive scores indicate peptides in the top 1% of the predictions while negative scores indicate peptides in the low 99% of the predictions for a single allele.

### Incorporating multiple specificity in HLA-I ligand predictors

All peptides annotated to the same allele were re-analyzed with our mixture model to identify multiple motifs. Peptides of different lengths were treated separately, always setting the core length equal to the peptide length in MixMHCp. Treating separately peptides of different lengths is important for multiple specificity analysis since peptides of different length may not display the same multiple specificity patterns (see example of C-terminal extensions in Fig. 4). All cases were then manually reviewed to determine whether the multiple motifs differed significantly from the single motif (Fig. 4).

**FIGURE 4:**
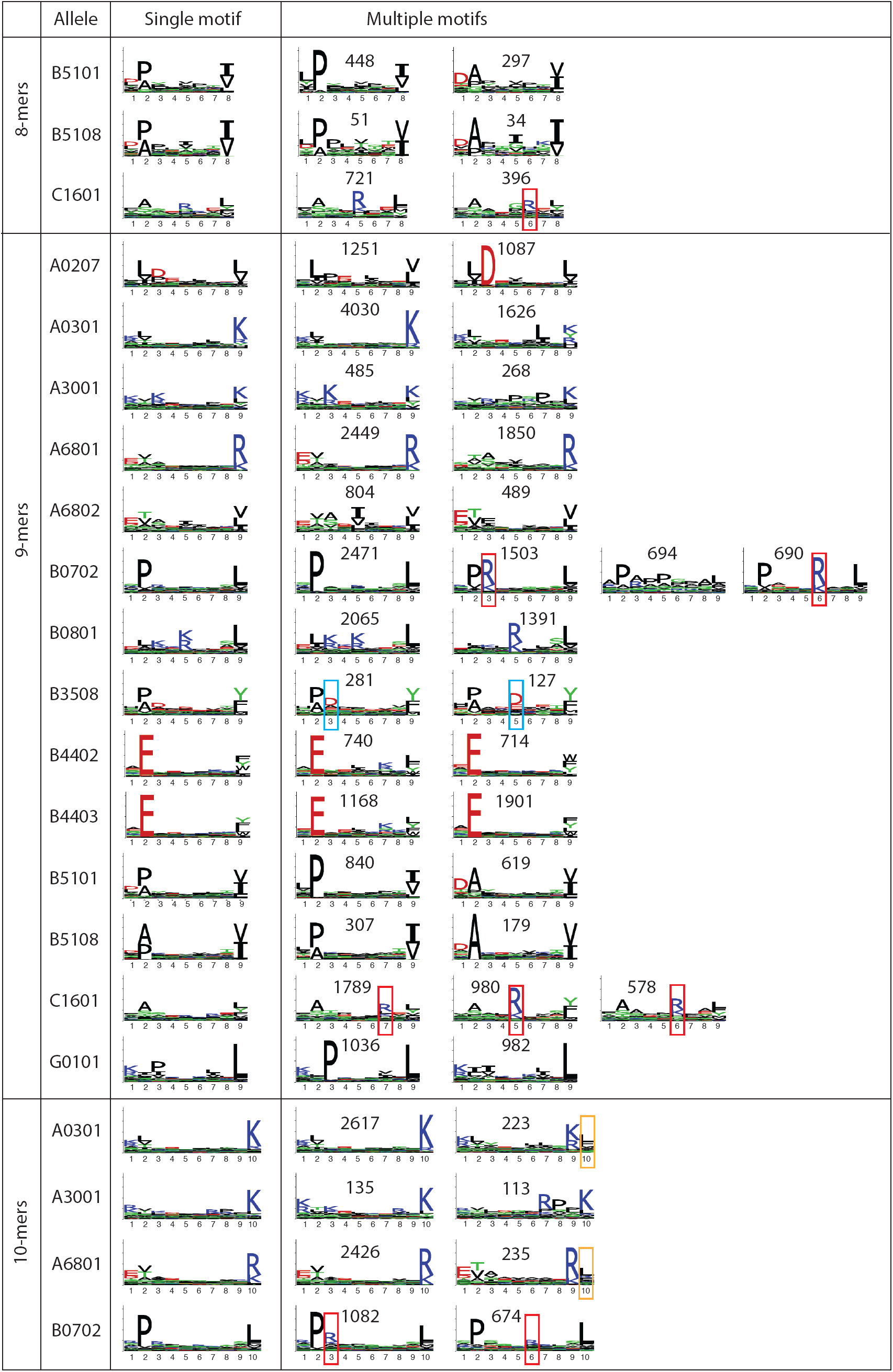
Cases of multiple specificity identified in this work for different alleles and different peptide lengths. The number of peptides supporting each motif is indicated. The red, respectively cyan, rectangles highlight cases of mutual exclusivity of positively, respectively negatively, charged residues. The orange rectangles highlight C-terminal extensions. The y-axis scale is the same for all logos (max at log_2_(20)).

To investigate whether multiple specificity could improve HLA-I ligand predictions, we modeled these cases with multiple PWMs using the framework introduced previously by ourselves for peptide recognition domains (19, 22) and recently applied to model C-terminal extensions in 10-mer HLA-I ligands (9). Peptides were assigned to each motif identified by the mixture model based on the highest responsibility values and multiple PWMs (*M*^(*h,L,k*)^) were built as described above for each allele *h*, each group of peptides *k* and each length *L*. Each motif also includes a weight *W*^*k*^ corresponding to the fraction of peptides assigned to it in the training set. The score of a new peptide of length *L* with allele *h* is then computed as:

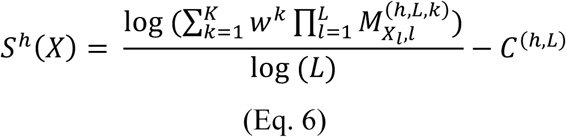

The new version of the HLA-I ligand predictor (MixMHCpred2.0.1) is available at https://github.com/GfellerLab/MixMHCpred.

### HLA peptidome profiling of meningioma samples with mass spectrometry based immunopeptidomics

Snap frozen meningioma tissues from patients were obtained from the Centre hospitalier universitaire vaudois (CHUV, Lausanne, Switzerland). HLA-I typing was performed for all samples. Informed consent of the participants was obtained following requirements of the institutional review board (Ethics Commission, CHUV). Protocol F-25/99 has been approved by the local Ethics committee and the biobank of the Lab of Brain Tumor Biology and Genetics.

We extracted HLA-I peptides from the meningioma samples as previously described (1). Briefly, snap-frozen meningioma tissue samples were placed in tubes containing ice cold PBS lysis buffer comprising of 0.25% sodium deoxycholate (Sigma-Aldrich), 0.2 mM iodoacetamide (Sigma-Aldrich), 1 mM EDTA, 1:200 Protease Inhibitors Cocktail (Sigma-Aldrich), 1 mM Phenylmethylsulfonylfluoride (Roche, Basel, Switzerland), 1% octyl-beta-D glucopyranoside (Sigma-Alrich) and homogenized on ice in 3-5 short intervals of 5 seconds each using an Ultra Turrax homogenizer (IKA, Staufen, Germany) at maximum speed. Cell lysis was performed at 4°C for 1 hour. Lysates were cleared by centrifugation at 20,000 rpm in a high speed centrifuge (Beckman Coulter, JSS15314, Nyon, Switzerland) at 4°C for 50 minutes. The lysates were loaded on stacked 96-well single-use micro-plates (3µm glass fiber, 10µm polypropylene membranes; ref number 360063, Seahorse Bioscience, North Billerica, Massachusetts). The first plate contained protein-A sepharose 4B (Pro-A) beads (Invitrogen, Carlsbad, California) for depletion of antibodies, and the second plate contained same beads cross-linked to W6/32 anti HLA monoclonal antibodies as previously described (1). We then employed the Waters Positive Pressure-96 Processor (Waters, Milford, Massachusetts). We washed the second plate with 4 times 2 mL of 150 mM sodium chloride (NaCl) (Carlo-Erba, Val de Reuil, France) in 20 mM Tris-HCl pH 8, 4 times 2 mL of 400 mM NaCl in 20 mM Tris-HCl pH 8 and again with 4 times 2 mL of 150 mM NaCl in 20 mM Tris-HCl pH 8. Finally, we washed the beads twice with 2 mL of 20 mM Tris-HCl pH 8. We stacked the affinity plate on top of a Sep-Pak tC_18_ 100 mg Sorbent 96-well plate (ref number: 186002321, Waters) already equilibrated with 1 mL of 80% ACN in 0.1 % TFA and with 2 mL of 0.1% TFA. We eluted the HLA and peptides with 500 µL 1% TFA into the Sep-Pak plate and then we washed this plate with 2 mL of 0.1 % TFA. Thereafter, we eluted the HLA-I peptides with 500 µL of 28% ACN in 0.1% TFA into a collection plate. Recovered HLA-I peptides were dried using vacuum centrifugation (Concentrator plus Eppendorf) and stored at −20°C.

Prior to MS analysis HLA-I peptide samples were re-suspended in 10 µL of 2% ACN/0.1 % FA and aliquots of 3 ul for each MS run were placed in the Ultra HPLC autosampler. All samples were acquired using the nanoflow Ultra-HPLC Easy nLC 1200 (Thermo Fisher Scientific, LC140) coupled online to a Q Exactive HF Orbitrap mass spectrometer (Thermo Fischer Scientific) with a nanoelectrospray ion source (Sonation, PRSO-V1, Baden-Württemberg, Germany) as previously described (1). We packed the uncoated PicoTip 8µm tip opening with 75 µm i.d. x 50 cm long analytical columns with ReproSil-Pur C18 (1.9 µm particles, 120 Å pore size, Dr. Maisch GmbH, Ammerbuch, Germany). Mounted analytical columns were kept at 50°C using a column oven. Data was acquired with data-dependent “top10” method. The MS scan range was set to 300 to 1,650 *m/z* with a resolution of 60,000 (200 *m/z*) and an AGC target value of 3e6 ions. For MS/MS, AGC target values of 1e5 were used with a maximum injection time of 120 ms at set resolution of 15,000 (200 *m/z*). In case of assigned precursor ion charge states of four and above, no fragmentation was performed.

We employed the MaxQuant computational proteomics platform version 1.5.5.1 (34) to search the peak lists against the UniProt databases (Human 42,148 entries, March 2017) and a file containing 247 frequently observed contaminants. We used default settings unless otherwise mentioned. Methionine oxidation (15.994915 Da), cysteine carbamidomethylation (57.021463 Da) and cysteinylation (119.004099) were set as variable modifications. The second peptide identification option in Andromeda was enabled. A false discovery rate (FDR) of 0.01 and no protein FDR was set. The enzyme specificity was set as unspecific and the ‘match between runs’ option was enabled across different replicates of the same biological sample in a time window of 0.5 min and an initial alignment time window of 20 min. We used the ‘peptides’ MaxQuant output table from which peptides matching to reverse and contaminants were filtered out (Supplemental Table III). MS RAW data, MaxQuant parameters and selected MaxQuant output tables used for analyses have been deposited to the ProteomeXchange Consortium (35) via the PRIDE partner repository with the dataset identifier PXD009925.

Two of these samples (3911-ME_I and 4021_I) were included in the peptide length distribution and multiple specificity analysis as well as the training of the new version of the HLA-I ligand predictor (MixMHCpred2.0.1), since they contained alleles (HLA-B49:01 and HLA-C15:05) whose motifs were not detected in our collection of publicly available HLA peptidomics studies, and therefore enabled us to expand the allelic coverage of MixMHCpred. The other ten datasets were kept for benchmarking of the new version of MixMHCpred (Fig. 6) and were not included in any training procedures presented in this work.

### Benchmarking MixMHCpred2.0.1

To benchmark our new algorithm (MixMHCpred2.0.1), we used the ten HLA peptidomes from meningioma samples that were measured in this study. Negative data were chosen by randomly selecting peptides from the human proteome so as to have 4 times more negative than positive. Peptides of lengths 8 to 14 were uniformly selected in the negative data. When attempting to predict peptides naturally presented in meningioma samples with many (i.e. up to six) different HLA-I alleles, we used for each peptide the highest score of MixMHCpred2.0.1 across all alleles. Precision in the top 20% of the predicted peptides (which is also equal to recall in the case of 4-fold excess of negative data) and Area Under the ROC Curve (AUC) were computed for each sample (4^th^ bars in Fig. 6A And 6B). AUC values were also computed with 99-fold excess of random negative data (Fig. 6C). MixMHCpred2.0.1 scores were then computed in the absence of peptide length distribution corrections and multiple specificity (1^st^ bars in Fig. 6), absence of peptide length distribution corrections (2^nd^ bars in Fig. 6) or absence of multiple specificity (3^rd^ bar in Fig. 6). When benchmarking with NetMHCpan4.0 (17), we used the percentile rank of the elution scores and chose the lowest percentile rank across all alleles for each peptide (5^th^ bar in Fig. 6).

### X-ray structures of HLA-I alleles

X-ray structures of HLA-I alleles shown in Fig. 2A were retrieved from the PDB. A representative structure for each allele was selected based on (i) the fact that the ligand is a 9-mer (ii) the absence of other receptors binding to the peptide-HLA-β2m complex (e.g., TCR or KIR, the only exception is HLA-C03:04 for which a crystal structure was only available in complex with KIR2DL2) and (iii) the lowest R value. This led to a total of 30 alleles with X-ray structures in complex with 9-mers (Supplemental Table IV).

Euclidean distances between the peptides and the beta sheet of the HLA-I molecules were computed as the smallest distance of any heavy atom of the peptide (P3 – P Ω-1) with any heavy atom of residues in the beta sheet (positions 5, 6, 7, 8, 9, 10, 11, 12, 23, 24, 25, 26, 27, 28, 95, 96, 97, 98, 99, 100, 101, 113, 114, 115, 116, 117 and 118). Most of the time, the residue closest to the peptide in the beta sheet is residue 97, which is located in the center of the binding site and points towards the non-anchor residues of the ligand. In each structure, B factors for all Cα atoms of the peptides (P3 – P Ω −1) were averaged and this value was normalized (i.e. divided) by the average of B factors of all Cα atoms of the HLA-I molecule to enable meaningful comparison between different structures.

### In vitro refolding assays

All peptides used in refolding assays with HLA-A03:01, HLA-B07:02, HLA-B15:01 and HLA-C06:02 were synthesized with free N and C-termini (1mg of each peptide, > 80% purity). Peptides were incubated separately with denaturated HLA-I alleles refolded by dilution in the presence of biotinylated beta-2 microglobulin proteins at temperature T=4°C for 48 hours. The solution was then incubated at 37°C. Samples were retrieved at time t=0h, 8h, 24h, 48h and t=72h. Stable complexes indicating interactions between HLA-I molecules and the peptides were detected by ELISA. Two independent replicates were performed for each measurement in Fig. 2C, 5B and 5C. Positive controls were used to renormalize the ELISA absorbance values between the different replicates and the y-axis shows the mean and standard deviation of these normalized values. Values in Fig. 2C were further renormalized by the 9-mer at time t=0h.

## Results

### Peptide length distributions in HLA-I ligands

To study the length distribution of naturally presented HLA-I ligands, we used our collection of curated HLA peptidomics datasets, including two novel meningioma samples (4’719 unique 8- to 14-mer peptides, see Materials and Methods and Supplemental Table III), to reach a total of 227’510 unique peptides. We used our recent motif deconvolution algorithm MixMHCp (3) to assign allelic restriction in pooled HLA peptidomics samples and expanded this approach better handle peptides of different lengths and allow for a flat motif that can identify contaminants that do not fit any of the motifs learned by the algorithm, akin to the trash cluster in GibbsCluster (5, 6, 24) (Eq. 1-3 and Materials and Methods). Supplemental Table I shows the number of predicted contaminants for each sample analyzed in this work and each peptide length. Overall, we observed a median 3.7% of predicted contaminants across samples, with some samples displaying over 25% of predicted contaminants (e.g., HLA-A68:02 in (7), as also reported in (6)). The highest fraction of predicted contaminants was observed in samples obtained by transfecting soluble HLA-I alleles, followed by mono-allelic samples, and then pooled HLA peptidomics samples (Fig. 1A). We also observed a much higher fraction of predicted contaminants among longer peptides (median of 58.8% among 14-mers versus 0.66% among 9-mers, see Fig. 1B).

Focusing on alleles with enough MS data (>200 ligands after removing peptides assigned to the flat motif), we could estimate peptide length distributions for a total of 84 alleles (Supplemental Fig. 1). As previously reported (7, 16–18), we observed significant differences in peptide length distributions between different alleles. To have a global view of the diversity of peptide length distributions across HLA-I alleles, we used t-SNE to represent in 2D the different HLA-I alleles based on the similarity of their peptide length distributions and proposed three distinct clusters of HLA-I alleles (Fig. 2A and 2B) (see Materials and Methods). Cluster 1 is composed of HLA-I alleles that accommodate many longer ligands (e.g., HLA-A01:01 or HLA-03:01). Cluster 2 contains alleles that can bind 10- and 11-mers, in addition to canonical 9-mers, but much less longer peptides (e.g., HLA-A02:01). Finally, cluster 3 contains alleles with peptide length distributions peaked on 9-mers (e.g., HLA-C06:02) or 8- and 9-mers (e.g., HLA-B51:01) and much fewer longer peptides.

Some patterns emerged from this analysis in terms of which HLA-I alleles fall into which cluster. First, we observed a strong clustering of HLA-C alleles (mainly in cluster 3), whereas HLA-A and HLA-B alleles were more mixed, with few HLA-A alleles in cluster 3. As expected, HLA-B alleles found in cluster 3 also displayed peptide length distributions peaked at 8- and 9-mers and include alleles with anchor residues at P5 (e.g., HLA-B08:01, HLA-B14:02, HLA-B37:01) (3). To test whether the peptide length distributions peaked at 9-mers for HLA-C alleles reflected differences in binding affinity, we tested six different ligands of HLA-C06:02 (Materials and Methods and Fig. 2C). In all cases we observed lower stability of both the 10- and 11-mers (orange and cyan, respectively) compared to the 9-mers (purple). These results suggest that the preference of HLA-C06:02 for 9-mers compared to 10- or 11-mers is also due to lower binding affinity of longer peptides, in addition to the expected higher frequency of 9-mer peptides available for loading in the ER (16) Reversely, 8-mers (black) displayed only slightly lower stability compared to 9-mers, suggesting that their lower frequency in MS data is mainly due to lower frequency of 8-mers in the ER (16). Similar results were previously observed for HLA-B51:01 (16).

To explore the structural basis of peptide length distributions, we surveyed existing X-ray structures of HLA-I alleles in complex with 9-mer peptides (Supplemental Table IV) and measured the distance between the ligand and the beta sheet of the HLA-I binding site (see Materials and Methods). Interestingly, we observed that the smallest distance between residues in the beta sheet of the HLA-I binding site and residues at non-anchor positions (P3 to P8) in the peptide is significantly lower for alleles in cluster 2 and 3 compared to alleles in cluster 1 (Fig. 2D, see example in Fig. 2E). In addition, the average normalized B-factor of C*α* atoms of residues P3 to P8 (see Materials and Methods), which represents how flexible these atoms are in X-ray structures, is significantly lower for alleles found in cluster 2 and 3 compared to alleles in cluster 1 (Fig. 2F). The lower distance between the residues in the beta sheet and those in the peptide together with the lower B-factors of peptides for alleles in cluster 2 and 3 indicate that 9-mer ligands of these alleles are more constrained and less floppy. This provides a potential structural explanation for the observed differences in peptide length distributions between different alleles and suggests that bulging out of the peptides in the presence of additional amino acids (i.e., longer peptides) is less favorable in alleles from clusters 2 and 3.

To further support this hypothesis, we note that HLA-B57:01 and HLA-B58:01 differ by only two residues in the HLA-I binding site (47 and 97) and have very similar binding motifs (Fig. 2E). Position 97 is located close to non-anchor residues of the peptide (P3 to P8), while position 45 is buried in the B pocket (Fig. 2E). In HLA-B57:01 valine is found at position 97 and does not interact directly with residues P3-P8 in the peptide (PDB: 2RFX (36), minimal distance equal to 5.4Å), while in HLA-B58:01 arginine is found at position 97 and makes direct interactions with the peptide (PDB: 5IND (37), minimal distance equal to 3.5Å) (Fig. 2E). Consistent with the mechanism proposed above, HLA-B57:01 has a broader peptide length distribution than HLA-B58:01 (Fig. 2A). These data suggest that position 97 in the HLA-I binding site may play a role in determining length preference and Fig. 2B indicates that, on average, smaller residues are found in alleles of cluster 1. This is consistent with the increased interactions previously observed between W97 and the peptide in HLA-B14:02 compared to most B27 alleles which have N at position 97 (38).

### Predicting peptide length distributions

The clustering observed in Fig. 2A suggests that it may be possible to predict peptide length distribution for a new HLA-I allele based on its sequence only, by predicting to which cluster it belongs and then mapping the average peptide length distribution of the cluster to this new allele. To this end, we trained a multinomial logistic regression, using as features all the HLA-I residues found in the binding site and as output the 3 clusters identified in Fig. 2A (Materials and Methods). Five-fold cross-validation was carried out, including both the clustering and the training of the multinomial logistic regression. The average correlation between observed and predicted peptide length distributions was 0.972. As a comparison, a random assignment of HLA-I alleles to the clusters during the cross-validation would result in an average correlation of 0.930. This clearly indicates that our approach has predictive power for peptide length distributions based on HLA-I sequences, beyond what is conserved across all HLA-I alleles (e.g., the higher frequency of 9-mers).

### Amino acid frequencies at non-anchor positions

We next took advantage of our large collection of unbiased HLA-I ligands of different lengths to investigate the changes in amino acid preferences at non-anchor positions (P5 to P Ω −2) for alleles displaying at least 20 ligands for all peptide lengths *L=8,…14* (13 alleles in total, see Materials and Methods). Fig. 3 shows the changes in amino acid frequencies with respect to peptide lengths. The first pattern that emerges from this analysis is the very clear increase in frequency of glycine in longer peptides. Glycine is known to destabilize secondary structures and induce protein disorder (39). We anticipate that the higher frequency of glycine in longer HLA-I ligands enables them to adopt bulging conformations more easily. We also observed increased frequency of negatively charged residues (aspartate and glutamate), serine and to a lower extent alanine. Glycine, serine and alanine have small sidechains, which confirms at a larger scale the observation made for HLA-C04:01 ligands (40). Disfavored amino acids in longer peptides include leucine, tryptophan, arginine, valine, phenylalanine, histidine and isoleucine. Many of them tend to induce structured conformations of polypeptides, further indicating that, on average, longer HLA-I ligands tend to be less structured at non-anchor positions. As noted in previous studies (4, 7), we also observed a very low frequency of cysteine at all peptide lengths, which is likely due to the fact that cysteine modifications are not included in standard MS spectral searches.

### Multiple specificity in HLA-I ligands

To study cases of multiple specificity, we applied the new version of the MixMHCp algorithm to each dataset of HLA-I ligands, treating separately peptides of different lengths (Materials and Methods). All cases were manually reviewed and our results revealed several instances of multiple specificity (Fig. 4). These include the previously reported multiple specificity of HLA-B07:02 (3), as well as cases of C-terminal extensions in 10-mers for HLA-A03:01 and HLA-A68:01 (orange boxes in Fig. 4) (9). For HLA-B07:02, the multiple specificity predicts mutual exclusivity of positively charged residues at P3 and P6. In our previous work, we had proposed a structural explanation for this observation, involving the presence of aspartate in the HLA binding site at position 114 which can interact with one, but not two positively charged residues in the peptide (3). A similar multiple specificity pattern is observed for HLA-C16:01 (Fig. 4, red boxes), and this allele has also aspartate at position 114. For HLA-B35:08, existing structures indicate that R156 can interact with residues at P3 (PDB: 2AXF (41)) or P5 (PDB: 2FZ3 (42)) (Fig. 5A), which is consistent with the mutual exclusivity of negatively charged residues observed at these 2 positions in MS data (Fig. 4, cyan boxes).

**FIGURE 5:**
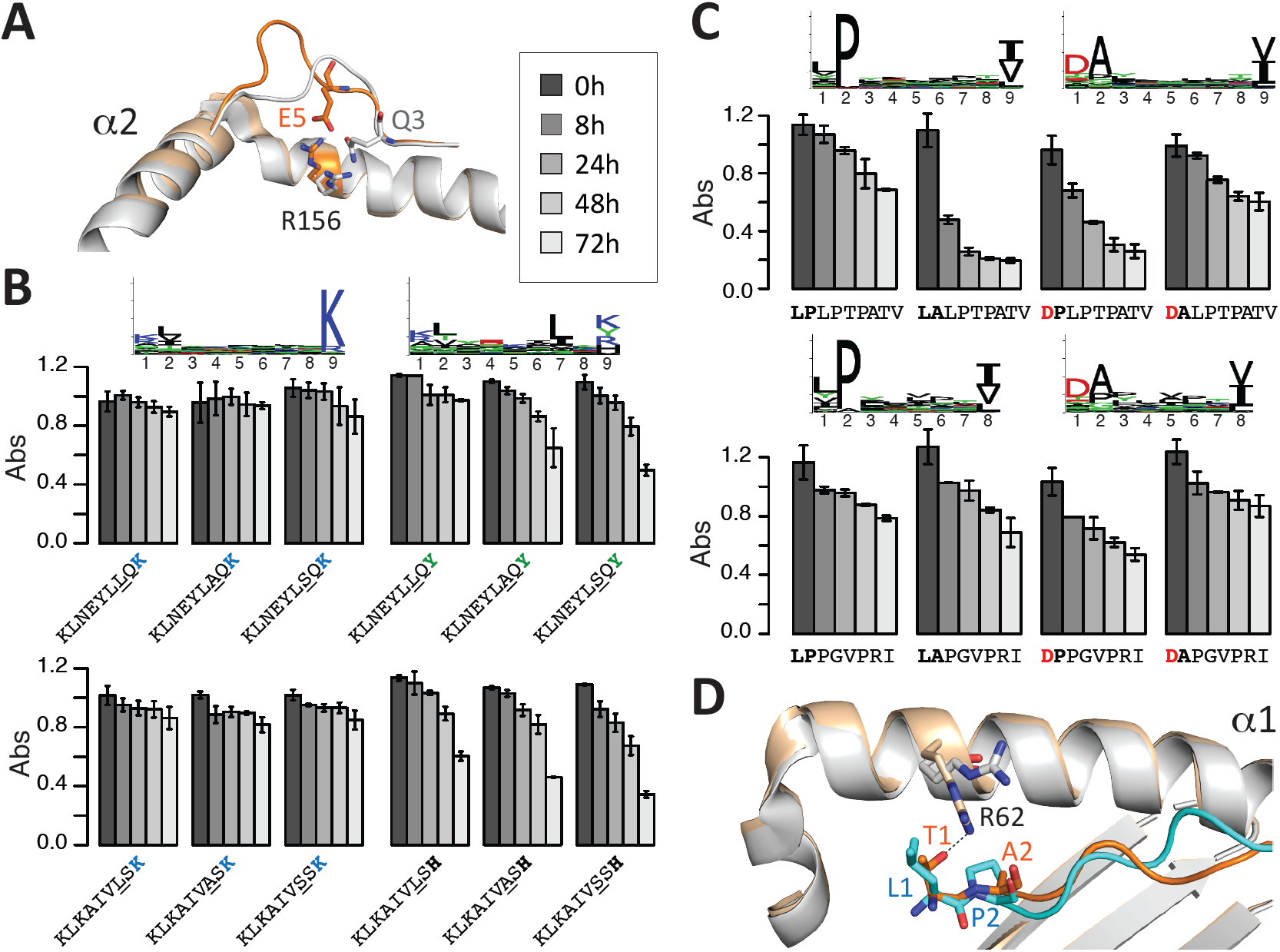
**A:** Superposition of X-ray structures of HLA-B35:08, indicating that R156 can interact with amino acids at P3 (PDB:2FZ3, grey) and P5 (PDB:2AXF, orange), consistent with the mutual exclusivity of negatively charged amino acids observed at these two positions in MS data (Fig. 4, cyan boxes). **B:** Validation of HLA-A03:01 multiple specificity: peptides with (K) at P9 do not show some specificity at P7, while peptides with Y at P9 do show specificity for L at P7. **C:** Validation of HLA-B51:01 multiple specificity: peptides with L_1_P_2_ or D_1_A_2_ display higher stability than peptides with L_1_A_2_ or D_1_P_2_. **D:** Superposition of X-ray structures of HLA-B51:01 in complex with two peptides matching the two motifs identified in MS data: ligand L_1_P_2_ in blue and HLA-B51:01 in grey (PDB:1E27); ligand T_1_A_2_ in orange and HLA-B51:01 in pale orange (PDB:1E28).

For two alleles (HLA-A03:01 and HLA-B51:01), we set out to validate these predictions. For HLA-A03:01, the multiple specificity model predicted that the binding of peptides with R/K at P9 should not be much influenced by the amino acid at P7, while peptides with Y/H at P9 should show additional specificity for leucine at P7. These predictions could be validated by observing that L7A mutation of peptides with K at P9 did not affect the binding, while L7A mutation decreased the binding of peptides with Y or H at P9 (Fig. 5B). For HLA-B51:01, peptides with sequences predicted to follow the two binding specificities (i.e., L_1_P_2_ and D_1_A_2_), as well as the other possibilities (i.e., D_1_P_2_ and L_1_A_2_) were tested. The latter are predicted to display weaker binding, although the single 9-mer logo of HLA-B51:01 would naively predict D_1_P_2_ to show the highest stability (Fig. 4). For both 9-mers and 8-mers, we could confirm the predictions of the multiple specificity analysis by observing that peptides with either L_1_P_2_ or D_1_A_2_ displayed higher stability than ligands with D_1_P_2_ or L_1_A_2_ (Fig. 5C). Comparison of HLA-B51:01 crystal structures in complex with two different ligands (L_1_P_2_ in blue and T_1_A_2_ in orange in Fig. 5D) (43) indicates that the polar group of T_1_ interacts with R62, which is consistent with the specificity for negatively charged or polar residues seen in MS data at P1. Reversely, in the presence of L_1_, the sidechain of R62 points towards the solvent. In addition, superposition of the two structures suggest that proline at P2 may not be compatible with the position of the threonine sidechain at P1. This suggests a possible explanation for the multiple specificity of HLA-B51:01. If alanine is found at P2, then negatively charged or polar amino acids are preferred at P1 and can make interactions with R62. Reversely, if proline is found at P2, amino acids at P1 cannot make direct interactions with R62, which is then pointing towards the solvent, and hydrophobic sidechains are preferred at P1. It is also interesting to note that one of the first neo-antigens identified by MS (DANSFLQSV) in melanoma samples (44) matches very precisely the second motif of HLA-B51:01 identified in this work.

### Incorporating peptide length distribution and multiple specificity in HLA-I ligand predictors

To incorporate peptide length distribution and multiple specificity in HLA-I ligand predictors, and explore whether this would lead to improved predictions of naturally presented HLA-I ligands, we developed a new version of MixMHCpred (v2.0.1). Peptide length distribution information was incorporated by adding a penalty so that the length distribution of the top 1% of peptides predicted from a pool of 700’000 random peptides (100’000 of each length *L=8,…,14*) follows by construction the distribution observed in MS data (Eq. 5 and Materials and Methods). For all alleles displaying multiple specificity, their binding specificity was modeled with multiple Position Weight Matrices (PWMs) (Eq. 6 and Materials and Methods).

To validate this new version of our predictor in an independent dataset, we measured the HLA peptidome of 10 new meningioma samples, resulting in 27,882 unique peptides of length 8 to 14 (Materials and Methods and Supplemental Table III). We then attempted to predict these peptides by adding 4 times more negative data consisting of 8- to 14-mer peptides randomly sampled from the human proteome (Materials and Methods). Fig. 6 shows the results in terms of precision in the top 20% of the predictions (which is equal to recall since we have a 4-fold excess of negative data) and Area Under the receiver operating Curve (AUC) values. In all cases, we saw clear improvement by adding correction for peptide length distributions (compare 1^st^ and 3^rd^ bars, as well as 2^nd^ and 4^th^ bars in Fig. 6A and 6B). Moreover, in all but two samples containing alleles that were shown to display multiple specificity (stars in Fig. 6), we observed that including multiple specificity led to equal or better results (compare 1^st^ and 2^nd^ bars, as well as 3^rd^ and 4^th^ bars in Fig. 6). When comparing with the other predictor of naturally presented HLA-I ligands that includes MS data in its training (i.e., NetMHCpan4.0 (17)), we could observe better performance in precision for all samples and in AUC values for all but one sample (compare 4^th^ and 5^th^ bars in Fig. 6). We repeated the AUC analysis in the presence of 99-fold excess of randomly selected negatives (Fig. 6C), and observed that here as well our predictor displayed higher accuracy compared to other predictors trained on MS data.

**FIGURE 6:**
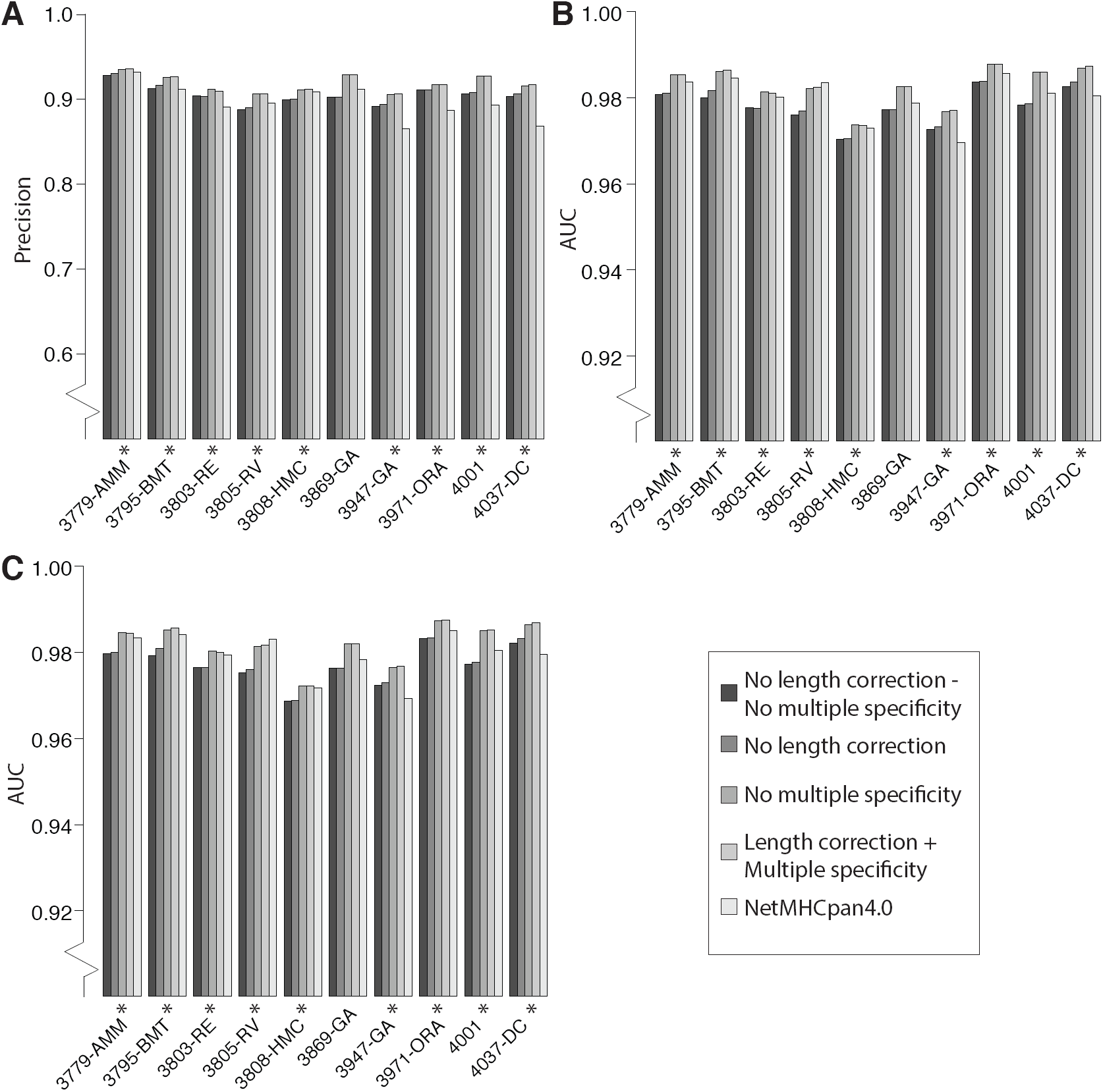
Improved predictions of naturally presented HLA-I ligands in 10 newly generated meningioma samples by explicitly modeling peptide length distribution and multiple specificity in MixMHCpred. **A:** Precision observed in the top 20% of the predicted peptides in the presence of 4-fold excess of randomly generated negative data (equivalent to recall). **B:** AUC values in the presence of 4-fold excess of randomly generated negative data. **C:** AUC values in the presence of 99-fold excess of randomly generated negatives. Stars indicate samples with at least one allele displaying multiple specificity.

## Discussion

In-depth and unbiased sampling of naturally presented HLA-I ligands is a powerful approach to unravel new properties of HLA-I molecules (3, 4, 6–9, 16, 40). In this work, we capitalized on these data to explore peptide length distribution and multiple specificity of HLA-I molecules at a much larger scale than previous studies. Our work revealed clear clustering of HLA-I alleles based on their peptide length distribution, which could guide investigation of the structural properties underlying the preference for peptides of different lengths. One clear pattern that emerged from our analysis is the restriction of HLA-C alleles to mainly 8- and 9-mers (8), likely reflecting both the biased distribution towards 9-mers of peptides in the ER (16) and the lower binding stability observed in Fig. 2C. Reversely, the similar binding stability observed between 8- and 9-mers suggests that the lower frequency of 8-mers among HLA-C naturally presented ligands is primarily due to their lower frequency in the pool of peptides available for loading in the ER (16), at least for HLA-C06:02. The restriction of HLA-C alleles to short peptides may also have functional implications. In addition to T Cell Receptors (TCRs) that can somatically rearrange their sequences to bind a wide range of HLA-I restricted targets, HLA-C alleles are the main targets of Natural Killer (NK) receptors which are germline encoded. Numerous crystal structures indicate the NK receptors bind to HLA-I alleles on top of the F pocket (12, 45). Longer peptides bulging out or extending at the C-terminus (9, 10, 13), may therefore interfere with the binding of NK receptors (12). This could explain why HLA-C alleles have evolved to preferentially bind to 8- and 9-mers.

Our work further enabled us to demonstrate, for the first time to our knowledge, that peptide length distributions can be predicted directly from HLA-I sequences, as shown in our cross-validation study.

The multiple specificity observed in this work suggests that, for some alleles, correlations between amino acid positions play an important role in binding to the HLA-I molecules (e.g., L_1_P_2_ and D_1_A_2_ being much more frequent to D_1_P_2_ or L_1_A_2_ in B51:01). These types of correlations, which we implicitly modelled with multiple PWMs, can also be captured with more complex machine learning frameworks, such as neural networks. It is worth noting that, from a mathematical point of view, multiple PWMs share many similarities with hidden nodes in neural networks, with the main difference being that the inputs of nodes in standard neural networks are combined with a logistic regression, whereas we use linear combinations in Eq. 6. As already observed for peptide recognition domains (19, 21, 22), an important advantage of the multiple PWMs framework is the straightforward visualization, which is useful to guide structural interpretation and experimental validation of the predictions. Nevertheless, the fact that only 10% of HLA-I alleles displayed clear multiple specificity and the modest improvement in prediction obtained by incorporating multiple specificity (Fig. 6) suggest that the assumption of positional independence is still a good first approximation for modeling the specificity of many HLA-I alleles (23).

Our performance estimates in Fig. 6 are only meant to compare between different methods and not to provide actual estimates of the success rate. The latter is unfortunately more difficult to assess in the absence of experimentally validated large negative datasets. In particular, in our benchmarking with 99-fold excess of randomly generated ‘negatives’ (Fig. 6C), we expect roughly 1% of them to be binders. This corresponds to the same number as the actual positives used in this benchmark and would make precision values in the top 1% more difficult to interpret.

Altogether, our work provides the first large-scale analysis of peptide length distributions and multiple specificity in naturally presented HLA-I ligands across a large panel of alleles. Including these two features into our predictor led to improved predictions of peptides presented on the surface of meningioma samples, and may therefore contribute to accelerating (neo-)antigen discovery in cancer immunotherapy.

## Acknowledgements

The computations were performed at the Vital-IT (http://www.vital-it.ch) Center for high-performance computing of the SIB Swiss Institute of Bioinformatics. We are thankful to Raphael Genolet for carrying out the HLA-I typing of the meningioma samples. We are thankful to Julien Schmidt and the Protein and Peptide Chemistry Facility from UNIL for synthesizing the peptides.

## Footnotes

This work was supported by the Swiss Cancer League (KFS-4104-02-2017-R), the Ludwig Institute for Cancer Research and by the ISREC Foundation thanks to a donation from the Biltema Foundation.

## Abbreviations

MS: Mass Spectrometry
ER: Endoplasmic Reticulum
PWM: Position Weight Matrix
AUC: Area Under the Curve
t-SNE: t- Distributed Stochastic Neighbor Embedding
IEDB: Immune Epitope DataBase
KIR: Killer-cell Immunoglobulin-like Receptor
PDB: Protein Data Bank
NK: Natural Killer

---

**Supplemental Figure 1:**
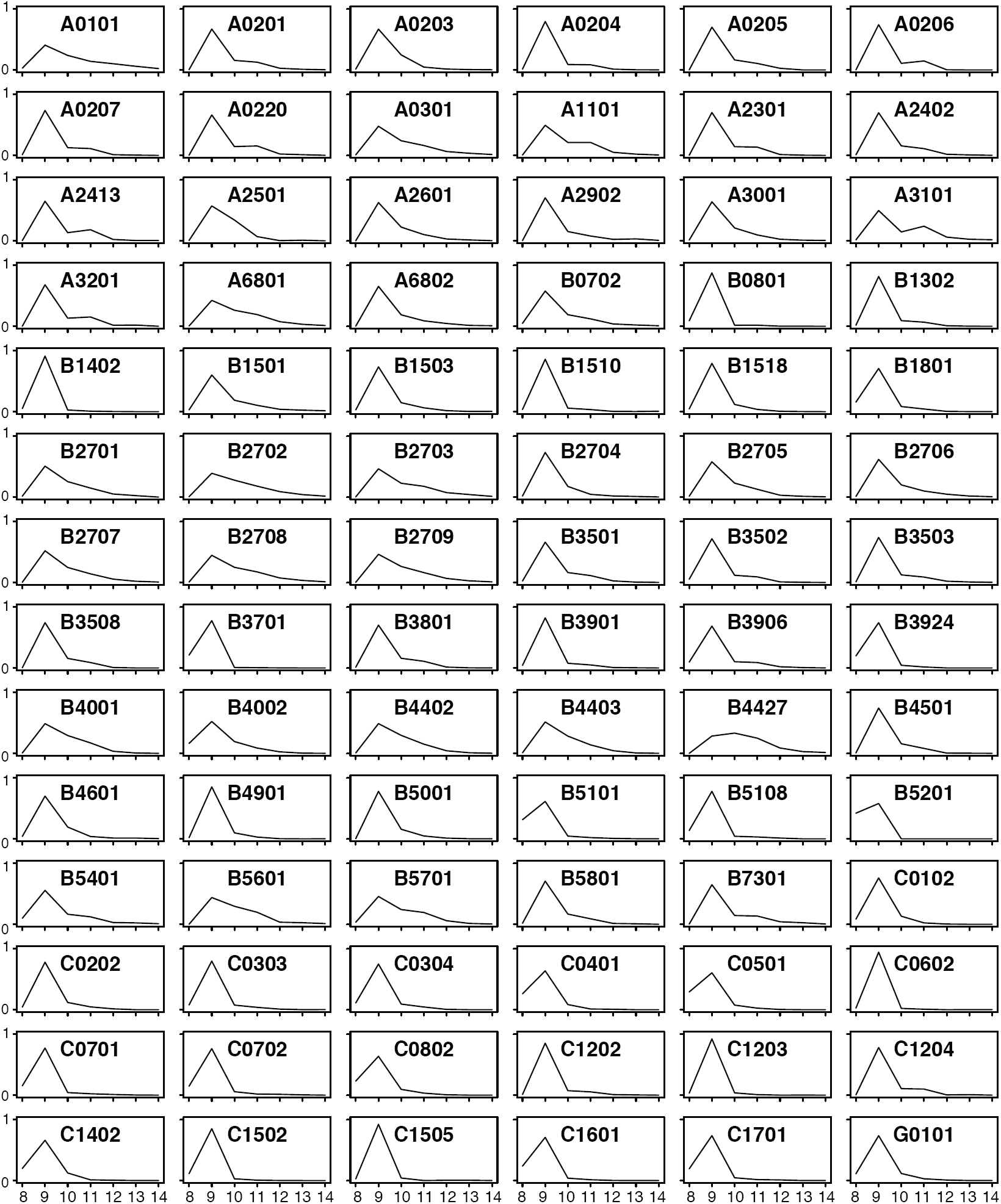
Peptide length distributions estimated from HLA peptidomics data for the 84 HLA-I alleles with >200 naturally presented ligands studied in this work.

## References

1 Bassani-Sternberg, M., S. Pletscher-Frankild, L. J. Jensen, and M. Mann. 2015. Mass spectrometry of human leukocyte antigen class I peptidomes reveals strong effects of protein abundance and turnover on antigen presentation. Mol. Cell Proteomics 14: 658–673.

2 Bassani-Sternberg, M., E. Bräunlein, R. Klar, T. Engleitner, P. Sinitcyn, S. Audehm, M. Straub, J. Weber, J. Slotta-Huspenina, K. Specht, M. E. Martignoni, A. Werner, R. Hein, D. H Busch, C. Peschel, R. Rad, J. Cox, M. Mann, and A. M. Krackhardt. 2016. Direct identification of clinically relevant neoepitopes presented on native human melanoma tissue by mass spectrometry. Nat Commun 7: 13404.

3 Bassani-Sternberg, M., and D. Gfeller. 2016. Unsupervised HLA Peptidome Deconvolution Improves Ligand Prediction Accuracy and Predicts Cooperative Effects in Peptide-HLA Interactions. J. Immunol.197: 2492–2499.

4 Bassani-Sternberg, M., C. Chong, P. Guillaume, M. Solleder, H. Pak, P. O. Gannon, L. E. Kandalaft, G. Coukos, and D. Gfeller. 2017. Deciphering HLA-I motifs across HLA peptidomes improves neo-antigen predictions and identifies allostery regulating HLA specificity. PLoS Comput. Biol.13: e1005725.

5 Andreatta, M., B. Alvarez, and M. Nielsen. 2017. GibbsCluster: unsupervised clustering and alignment of peptide sequences. Nucleic Acids Res.45: W458–W463.

6 Alvarez, B., C. Barra, M. Nielsen, and M. Andreatta. 2018. Computational Tools for the Identification and Interpretation of Sequence Motifs in Immunopeptidomes. Proteomics 18: e1700252.

7 Abelin, J. G., D. B. Keskin, S. Sarkizova, C. R. Hartigan, W. Zhang, J. Sidney, J. Stevens, W. Lane, G. L. Zhang, T. M. Eisenhaure, K. R. Clauser, N. Hacohen, M. S. Rooney, S. A. Carr, and C. J. Wu. 2017. Mass Spectrometry Profiling of HLA-Associated Peptidomes in Mono-allelic Cells Enables More Accurate Epitope Prediction. Immunity 46: 315–326.

8 Di Marco, M., H. Schuster, L. Backert, M. Ghosh, H.-G. Rammensee, and S. Stevanović. 2017. Unveiling the Peptide Motifs of HLA-C and HLA-G from Naturally Presented Peptides and Generation of Binding Prediction Matrices. J. Immunol.199: 2639–2651.

9 Guillaume, P., S. Picaud, P. Baumgaertner, N. Montandon, J. Schmidt, D. E. Speiser, G. Coukos, M. Bassani-Sternberg, P. Filippakopoulos, and D. Gfeller. 2018. The C-terminal extension landscape of naturally presented HLA-I ligands. Proc. Natl. Acad. Sci. U.S.A.115: 5083–5088.

10 McMurtrey, C., T. Trolle, T. Sansom, S. G. Remesh, T. Kaever, W. Bardet, K. Jackson, R. McLeod, A. Sette, M. Nielsen, D. M. Zajonc, I. J. Blader, B. Peters, and W. Hildebrand. 2016. Toxoplasma gondii peptide ligands open the gate of the HLA class I binding groove. Elife 5.

11 Nielsen, M., T. Connelley, and N. Ternette. 2017. Improved prediction of Bovine Leucocyte Antigens (BoLA) presented ligands by use of mass spectrometry-determined ligand-and in-vitro binding data. Journal of proteome research acs.jproteome.7b00675.

12 Pymm, P., P. T. Illing, S. H. Ramarathinam, G. M. O’Connor, V. A. Hughes, C. Hitchen, D. A. Price, B. K. Ho, D. W. McVicar, A. G. Brooks, A. W. Purcell, J. Rossjohn, and J. P. Vivian. 2017. MHC-I peptides get out of the groove and enable a novel mechanism of HIV-1 escape. Nature structural & molecular biology 219: 277.

13 Remesh, S. G., M. Andreatta, G. Ying, T. Kaever, M. Nielsen, C. McMurtrey, W. Hildebrand, B. Peters, and D. M. Zajonc. 2017. Unconventional Peptide Presentation by Major Histocompatibility Complex (MHC) Class I Allele HLA- A*02:01: BREAKING CONFINEMENT. J. Biol. Chem.292: 5262–5270.

14 Ritz, D., A. Gloger, B. Weide, C. Garbe, D. Neri, and T. Fugmann. 2016. High-sensitivity HLA class I peptidome analysis enables a precise definition of peptide motifs and the identification of peptides from cell lines and patients’ sera. Proteomics 16: 1570–1580.

15 Gfeller, D., and M. Bassani-Sternberg. 2018. Predicting Antigen Presentation-What Could We Learn From a Million Peptides? Front Immunol 9: 1716.

16 Trolle, T., C. P. McMurtrey, J. Sidney, W. Bardet, S. C. Osborn, T. Kaever, A. Sette, W. H. Hildebrand, M. Nielsen, and B. Peters. 2016. The Length Distribution of Class I-Restricted T Cell Epitopes Is Determined by Both Peptide Supply and MHC Allele-Specific Binding Preference. J. Immunol.196: 1480–1487.

17 Jurtz, V., S. Paul, M. Andreatta, P. Marcatili, B. Peters, and M. Nielsen. 2017. NetMHCpan-4.0: Improved Peptide-MHC Class I Interaction Predictions Integrating Eluted Ligand and Peptide Binding Affinity Data. J. Immunol.199: 3360–3368.

18 Andreatta, M., and M. Nielsen. 2016. Gapped sequence alignment using artificial neural networks: application to the MHC class I system. Bioinformatics 32: 511–517.

19 Gfeller, D., F. Butty, M. Wierzbicka, E. Verschueren, P. Vanhee, H. Huang, A. Ernst, N. Dar, I. Stagljar, L. Serrano, S. S. Sidhu, G. D. Bader, and P. M. Kim. 2011. The multiple-specificity landscape of modular peptide recognition domains. Mol. Syst. Biol.7: 484.

20 Nielsen, M., C. Lundegaard, P. Worning, S. L. Lauemøller, K. Lamberth, S. Buus, S. Brunak, and O. Lund. 2003. Reliable prediction of T-cell epitopes using neural networks with novel sequence representations. Protein Sci.12: 1007–1017.

21 Gfeller, D. 2012. Uncovering new aspects of protein interactions through analysis of specificity landscapes in peptide recognition domains. FEBS letters 586: 2764–2772.

22 Kim, T., M. S. Tyndel, H. Huang, S. S. Sidhu, G. D. Bader, D. Gfeller, and P. M. Kim. 2012. MUSI: an integrated system for identifying multiple specificity from very large peptide or nucleic acid data sets. Nucleic Acids Res.40: e47.

23 Peters, B., W. Tong, J. Sidney, A. Sette, and Z. Weng. 2003. Examining the independent binding assumption for binding of peptide epitopes to MHC-I molecules. Bioinformatics (Oxford, England) 19: 1765–1772.

24 Andreatta, M., O. Lund, and M. Nielsen. 2013. Simultaneous alignment and clustering of peptide data using a Gibbs sampling approach. Bioinformatics 29: 8–14.

25 Guasp, P., C. Alvarez-Navarro, P. Gomez-Molina, A. Martín-Esteban, M. Marcilla, E. Barnea, A. Admon, and J. A. López de Castro. 2016. The Peptidome of Behçet’s Disease-Associated HLA-B*51:01 Includes Two Subpeptidomes Differentially Shaped by Endoplasmic Reticulum Aminopeptidase 1. Arthritis & Rheumatology (Hoboken, N.J.) 68: 505–515.

26 Hilton, H. G., C. P. McMurtrey, A. S. Han, Z. Djaoud, L. A. Guethlein, J. H. Blokhuis, J. L. Pugh, A. Goyos, A. Horowitz, R. Buchli, K. W. Jackson, W. Bardet, D. A. Bushnell, P. J. Robinson, J. L. Mendoza, M. E. Birnbaum, M. Nielsen, K. C. Garcia, W. H. Hildebrand, and P. Parham. 2017. The Intergenic Recombinant HLA-B*46: 01 Has a Distinctive Peptidome that Includes KIR2DL3 Ligands. Cell Rep 19: 1394–1405.

27 Gloger, A., D. Ritz, T. Fugmann, and D. Neri. 2016. Mass spectrometric analysis of the HLA class I peptidome of melanoma cell lines as a promising tool for the identification of putative tumor-associated HLA epitopes. Cancer Immunol. Immunother.65: 1377–1393.

28 Mommen, G. P. M., C. K. Frese, H. D. Meiring, J. van Gaans-van den Brink, A. P. J. M. de Jong, C. A. C. M. van Els, and A. J. R. Heck. 2014. Expanding the detectable HLA peptide repertoire using electron-transfer/higher-energy collision dissociation (EThcD). Proc. Natl. Acad. Sci. U.S.A.111: 4507–4512.

29 Pearson, H., T. Daouda, D. P. Granados, C. Durette, E. Bonneil, M. Courcelles, A. Rodenbrock, J.-P. Laverdure, C. Côté, S. Mader, S. Lemieux, P. Thibault, and C. Perreault. 2016. MHC class I-associated peptides derive from selective regions of the human genome. J. Clin. Invest.126: 4690–4701.

30 Ritz, D., A. Gloger, D. Neri, and T. Fugmann. 2017. Purification of soluble HLA class I complexes from human serum or plasma deliver high quality immuno peptidomes required for biomarker discovery. Proteomics 17: 1600364.

31 Vita, R., J. A. Overton, J. A. Greenbaum, J. Ponomarenko, J. D. Clark, J. R. Cantrell, D. K. Wheeler, J. L. Gabbard, D. Hix, A. Sette, and B. Peters. 2015. The immune epitope database (IEDB) 3.0. Nucleic Acids Res.43: D405–412.

32 Van der Maaten, L., and G. E. Hinton. 2008. Visualizing High-Dimensional Data Using t-SNE. Journal of Machine learning Research 9: 2579.

33 Lund, O., M. Nielsen, C. Lundegaard, C. Keşmir, and S. Brunak. 2005. Immunological Bioinformatics, MIT Press.

34 Tyanova, S., T. Temu, and J. Cox. 2016. The MaxQuant computational platform for mass spectrometry-based shotgun proteomics. Nat Protoc 11: 2301–2319.

35 Vizcaíno, J. A., A. Csordas, N. Del-Toro, J. A. Dianes, J. Griss, I. Lavidas, G. Mayer, Y. Perez-Riverol, F. Reisinger, T. Ternent, Q.-W. Xu, R. Wang, and H. Hermjakob. 2016. 2016 update of the PRIDE database and its related tools. Nucleic acids research 44: 11033–11033.

36 Chessman, D., L. Kostenko, T. Lethborg, A. W. Purcell, N. A. Williamson, Z. Chen, L. Kjer-Nielsen, N. A. Mifsud, B. D. Tait, R. Holdsworth, C. A. Almeida, D. Nolan, W. A. Macdonald, J. K. Archbold, A. D. Kellerher, D. Marriott, S. Mallal, M. Bharadwaj, J. Rossjohn, and J. McCluskey. 2008. Human leukocyte antigen class I-restricted activation of CD8+ T cells provides the immunogenetic basis of a systemic drug hypersensitivity. Immunity 28: 822–832.

37 Li, X., P. A. Lamothe, R. Ng, S. Xu, M. Teng, B. D. Walker, and J.-H. Wang. 2016. Crystal structure of HLA-B*5801, a protective HLA allele for HIV-1 infection. Protein & cell 7: 761–765.

38 Kumar, P., A. Vahedi-Faridi, W. Saenger, E. Merino, J. A. López de Castro, B. Uchanska-Ziegler, and A. Ziegler. 2009. Structural basis for T cell alloreactivity among three HLA-B 14 and HLA-B 27 antigens. J. Biol. Chem.284: 29784–29797.

39 Linding, R., R. B. Russell, V. Neduva, and T. J. Gibson. 2003. GlobPlot: Exploring protein sequences for globularity and disorder. Nucleic acids research 31: 3701–3708.

40 Schittenhelm, R. B., N. L. Dudek, N. P. Croft, S. H. Ramarathinam, and A. W. Purcell. 2014. A comprehensive analysis of constitutive naturally processed and presented HLA-C*04:01 (Cw4)-specific peptides. Tissue antigens 83: 174–179.

41 Tynan, F. E., D. Elhassen, A. W. Purcell, J. M. Burrows, N. A. Borg, J. J. Miles, N. A. Williamson, K. J. Green, J. Tellam, L. Kjer-Nielsen, J. McCluskey, J. Rossjohn, and S. R. Burrows. 2005. The immunogenicity of a viral cytotoxic T cell epitope is controlled by its MHC-bound conformation. The Journal of experimental medicine 202: 1249–1260.

42 Miles, J. J., N. A. Borg, R. M. Brennan, F. E. Tynan, L. Kjer-Nielsen, S. L. Silins, M. J. Bell, J. M. Burrows, J. McCluskey, J. Rossjohn, and S. R. Burrows. 2006. TCR alpha genes direct MHC restriction in the potent human T cell response to a class I-bound viral epitope. Journal of immunology (Baltimore, Md.: 1950) 177: 6804–6814.

43 Maenaka, K., T. Maenaka, H. Tomiyama, M. Takiguchi, D. I. Stuart, and E. Y. Jones. 2000. Nonstandard peptide binding revealed by crystal structures of HLA- B* 5101 complexed with HIV immunodominant epitopes. J. Immunol.165: 3260–3267.

44 Kalaora, S., E. Barnea, E. Merhavi-Shoham, N. Qutob, J. K. Teer, N. Shimony, J. Schachter, S. A. Rosenberg, M. J. Besser, A. Admon, and Y. Samuels. 2016. Use of HLA peptidomics and whole exome sequencing to identify human immunogenic neo-antigens. Oncotarget 7: 5110–5117.

45 Boyington, J. C., S. A. Motyka, P. Schuck, A. G. Brooks, and P. D. Sun. 2000. Crystal structure of an NK cell immunoglobulin-like receptor in complex with its class I MHC ligand. Nature 405: 537–543.

